# Spatially resolved molecular analysis of host response to medical device implantation using the 3D OrbiSIMS highlights a critical role for lipids

**DOI:** 10.1101/2023.08.18.553860

**Authors:** Waraporn Suvannapruk, Leanne E Fisher, Jeni C Luckett, Max K Edney, Anna M Kotowska, Dong-Hyun Kim, David J Scurr, Amir M Ghaemmaghami, Morgan R Alexander

## Abstract

A key goal for implanted medical devices is that they do not elicit a detrimental immune response. Macrophages play critical roles in modulation of the host immune response and are the major cells responsible for persistent inflammatory reactions to implanted biomaterials. We investigate two novel immune-instructive polymers that stimulate pro- or anti-inflammatory responses from macrophages *in vitro*. These also modulate *in vivo* foreign body responses (FBR) when implanted subcutaneously in mice as coatings on biomedical grade silicone rubber. The tissue surrounding the implant is mechanically sectioned and imaged to assess the response of the polymers compared to silicone rubber. Immunofluorescent staining reveals responses consistent with pro- or anti-inflammatory responses previously described for these polymers. We apply 3D OrbiSIMS analysis to provide spatial analysis of the metabolite signature in the tissue surrounding the implant for the first time, providing molecular histology insight into the metabolite response in the host tissue. For the pro-inflammatory coating, monoacylglycerols (MG) and diacylglycerols (DG) are observed at increased intensity, while for the anti-inflammatory coating the number of phospholipid species detected decrease and pyridine and pyrimidine levels were elevated. These findings link to observations of small molecule signature from single cell studies of M2 macrophages *in vitro* where cell and tissue ion intensities were found to correlate suggesting potential for prediction. This illustrates the power of metabolite characterization by the 3D OrbiSIMS to gain insight into the mechanism of bio-instructive materials as medical devices and to inform on the FBR to biomaterials.

## 1. Introduction

Medical devices are ubiquitous in modern medicine, from coronary stents, catheters, hip/knee replacements, to the everyday contact lens. Patients can suffer adverse immune reactions to implanted devices, leading to chronic inflammation, pain, and on occasion, implant failure (1). The foreign body response (FBR) and chronic inflammation in the implant microenvironment can be detrimental to the function of implanted materials/tissues and increase patient mortality and morbidity (2). Macrophages play a critical role in orchestrating the FBR to implanted materials (3, 4), and can perpetuate chronic inflammation or enhance tissue healing depending on the phenotype they adopt in response to different biomaterials (5, 6). Therefore, control of inflammatory responses by modulating macrophage phenotype may improve implant integration. This has led to significant interest in designing novel non-eluting bio-instructive materials that interact positively with the immune system to induce a favorable macrophage response to medical devices (7–11).

The traditional approach to characterize macrophage phenotype during the FBR relies on immunohistochemistry for markers that are typically associated with pro-inflammatory and anti-inflammatory macrophages, such as nitric oxide synthase (iNOS) and the anti-inflammatory arginise-1 (Arg-1) respectively (12–14). One limitation of this approach is the co-expression of both iNOS and Arg-1 markers on many macrophages that makes it difficult to determine their functional phenotype accurately, and therefore a range of cell surface markers have also been identified by immunohistochemistry to profile macrophages (15). As an alternative to immunohistochemistry, here we investigate an approach using the metabolomic signature of cells and tissues to identify changes within the small molecule population at the host/material interface and provide insight into the related molecular changes within the tissue.

Metabolomics is defined as the comprehensive analysis of metabolites in a biological sample, and is a powerful technique that has the potential to improve diagnosis and treatment (16). Metabolomic information can provide an in-depth understanding of complex molecular interactions within biological systems and provide information that relates to cell phenotype (17). Technologies based on metabolomic markers have been established to study disease, here we explore its power for assessing the influence of bio-instructive implants.

Studies of metabolites in tissues have used a variety of instrumental and data processing techniques based on targeted and/or non-targeted techniques such as liquid chromatography-mass spectrometry (LC-MS), liquid extraction surface analysis-MS (LESA-MS), MALDI-imaging MS and secondary ion MS (SIMS). LC-MS is the most commonly used analytical method for detecting metabolites (18). Typically, LC-MS-based metabolite analyses proceed with the extraction of metabolites from tissue samples. It requires a significant amount of tissue, for example as presented by Woodward et al., using metabolomics to classify brain tumor tissue required at least 10 mg to obtain a sufficient signal (19). The sample is often prepared using solvent extraction of a tissue surface sample to isolate and inject to separate and then ionize analytes of interest (20).

A direct analysis alternative is LESA-MS, used by Meurs et al., to identify metabolites in paediatric ependymoma tumour tissue (21). LESA-MS has the advantages of sensitivity and high resolution mass analysers, but the limitations of these approaches are their low spatial resolution (500 μm-1 mm)(22). Time-of-flight secondary ion MS (ToF-SIMS) has also been used to identify small molecules in biological samples (23, 24). For application in animal studies, Palmquist et al., performed chemical and structural analysis of the bone-implant interface using ToF-SIMS (25). However, ToF-SIMS has been limited by its relatively low mass resolving power and thus has not been applied to metabolite identification due to associated limitations in assignment certainty (26). In the pursuit of greater assignment specificity of ions, a hybrid SIMS instrument with a high resolution Orbitrap mass analyser has been developed, termed 3D OrbiSIMS (27).

3D OrbiSIMS is a direct surface analysis technique which enables identification of biomolecules in complex samples using intact molecular ions, generated by an argon gas cluster primary ion beam with 2 µm spatial resolution, high mass resolving power (>240,000 for a peak of m/z 200), and excellent mass accuracy (<2 ppm) (27). This capability has been demonstrated on a variety of tissue and cell samples, assigning lipids, proteins, amino acids, peptide fragments of proteins and a selection of other small molecules (28–31). Furthermore, studies of metabolomics that use 3D OrbiSIMS do not need to use protocols necessary with other techniques including, chemically fixed cells, liquid extraction procedures, antibody-based cell markers or staining. Recently, 3D OrbiSIMS imaging has been used to observe the metabolite characteristics in brain tumor tissue samples and to identify clinically relevant molecular metabolism driven subgroup specific phenotypes, using a sample preparation approach similar to previous studies (21, 27).

Here, we applied 3D OrbiSIMS analysis to investigate whether there was a molecular signature for different pro- or anti-inflammatory macrophages at the interface of tissue implanted materials illustrated in Figure 1. We retrieved implants (silicone catheter segments) with or without immune-instructive polymer coatings described previously (x) 28 days after implantation in a subcutaneous murine model using freeze drying prior to analysis. Characteristic metabolites from the histological sections adjacent to the implanted foreign body site were identified using a combination of targeted database approaches to identifying lipids, and a data driven multivariate approach to highlight amino acids and other small molecules that we interpreted by comparison with literature on macrophage metabolomics.

**Figure 1.**
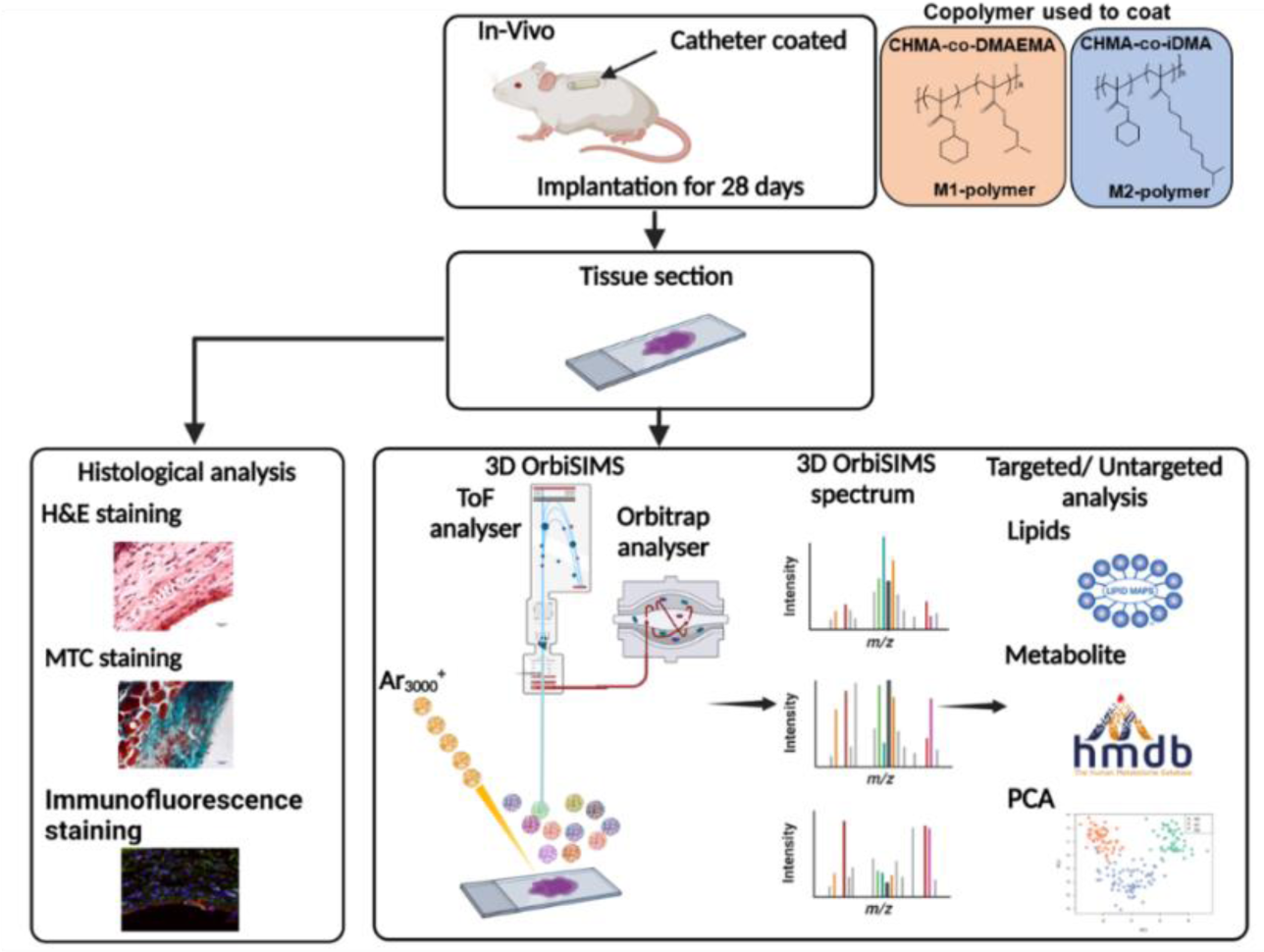
Schematic of the *in vivo* study experimental procedure. Catheters coated with copolymers were implanted subcutaneously into a mouse model of FBR for 28 days. Following implantation, fresh tissue samples were cut and mounted onto glass slides. For histological analysis, tissue sections were stained with H&E to assess tissue structure, MTC to analyse collagen thickness as an indication of fibrosis and IHC stains to characterise the macrophage phenotype at the catheter-tissue interface. For 3D OrbiSIMS, tissue slides were washed with ammonia formate, plunge frozen in liquid nitrogen and then freeze dried. The GCIB was rastered across the tissue section with the Orbitrap analyser collecting the high resolution mass spectrum, followed by multivariate analysis to undertake unbiased sample comparison, complemented with targeted analysis.

## 2. Results and discussion

### 2.1. Pro- and anti-inflammatory macrophage instructive polymers influence immune cell infiltration and collagen deposition

Polymers eliciting *in vitro* polarization of monocytes to M1 inducing pro-inflammatory macrophage phenotype or to M2 inducing anti-inflammatory macrophage phenotype were synthesized by thermal free radical polymerization and coated onto segments of clinical-grade silicone catheter via dipping. The polymers used had previously been identified to polarize macrophages *in vitro* and modulate FBR *in vivo*: M1 inducing: poly(cyclohexyl methacrylate-co-dimethylamino-ethyl methacrylate), abbreviated to pCHMA-DMAEMA induced a pro-inflammatory macrophage phenotype, and M2 inducing: poly(cyclohexyl methacrylate-co-isodecyl methacrylate), abbreviated to pCHMA-iDMA induced an anti-inflammatory macrophage phenotype (12). The *in vivo* responses to these novel polymer coatings were compared to each other and the widely employed biomedical polymer, silicone rubber, by subcutaneous implantation into a mouse model of FBR for 28 days. Upon recovery of the implants and surrounding skin, tissues were snap frozen in liquid nitrogen sectioned and freeze dried before 3D OrbiSIMS analysis. Adjacent sections were stained with H&E and MTC and immunohisto chemically labelled with iNOS and arginase 1 markers before optical microscopy to characterize the inflammatory response, collagen deposition and phenotype marker respectively (12). The staining and labelling revealed that the total number of macrophages was invariant between the different samples as seen in Figure S1A, Supporting Information but the anti-inflammatory coating did result in a far higher M2/M1 ratio of macrophages in the tissue near the catheter and the pro-inflammatory coating a far lower ratio with the uncoated PDMS implants in between (Figure S1D, Supporting Information).

A lower number of neutrophils were recruited to the M2-polymer (Figure S1B, Supporting Information), although this was not statistically significant. A lower number on the M2-polymer would be consistent with phagocytosis-induced cell death at the earlier stage of the inflammatory response. The thickness of collagen deposition at the surface of the anti-inflammatory polymer was significantly greater than the pro-inflammatory polymer and the uncoated catheters (Figure S1C, Supporting Information). These results are consistent with previous *in vivo* studies of these polymers undertaken in the same model (12).

### 2.2. Characterisation of metabolite changes in tissue local to implants

We analyzed the tissue areas surrounding the implant using 3D OrbiSIMS in both positive and negative secondary ion modes to provide molecular histology to complement the assignments provided by the immunohistochemistry. Principal component analysis (PCA) was initially applied as an untargeted data analysis approach to distinguish chemical differences in the complex 3D OrbiSIMS spectra from the local host-implant interface spectra. The *scores* plot of the first and second principal components is presented in Figure 2A, highlighting that there are chemical differences between the tissue proximal to the M2- and M1-inducing polymers and the uncoated silicone catheter. The *loadings* plot in Figure 2B presents the spectral components responsible for the chemical separation of the samples, in this case dominated by the glycerolipid fragments associated preferentially with the pro-inflammatory polymer implant. The positively loaded ions are assigned to glycerol lipids including monoacylglycerols (MG) and diacylglycerols (DG) at *m/z* 551.5035 (MG 32:2), *m/z* 575.5035 (MG 34:4), *m/z* 557.5191(MG 34:3) and *m/z* 601.5193 (DG O-36:5). The negative score on PC1 was associated with amino acids from the anti-inflammatory polymer highlighting a series of lower mass peaks, including m/z 91.0545 and *m/z* 130.0652 (tryptophan), and m/z 103.0543 and m/z 120.0808 (phenylalanine) suggesting higher protein levels in these samples, consistent with the greater collagen thickness formed in response to these implants (Figure S1C, Supporting Information).

**Figure 2.**
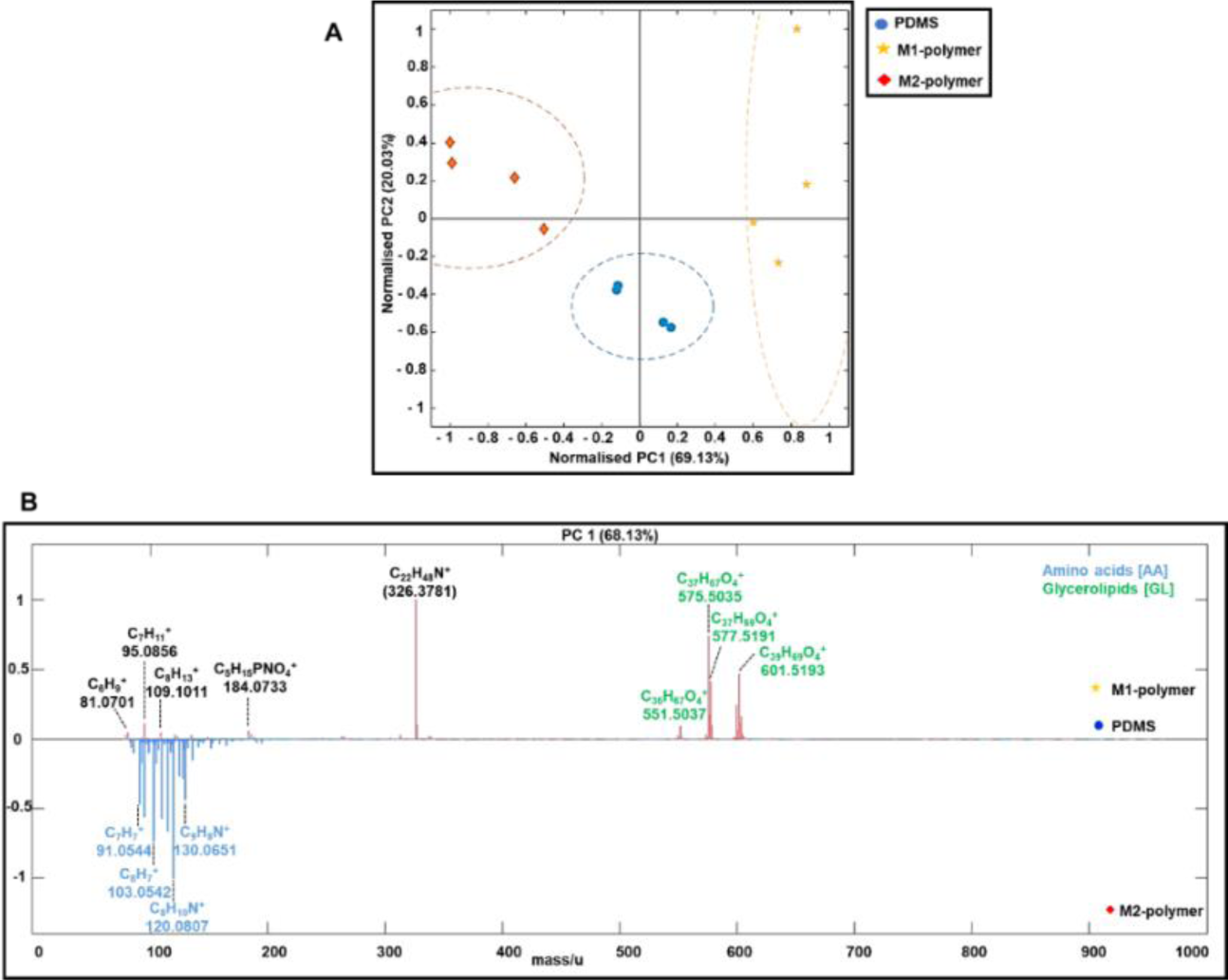
Principal component analysis (scores and loadings) for different tissue samples. (A) Principal component scores plot of PC1 and PC2 for the 3D OrbiSIMS spectra of PDMS, M1 polymer and M2 polymer tissue section samples on positive polarity. (B) Principal component analysis of 3 different tissue samples, loadings plots of the first (PC1) and (PC2) principal components and peak were assigned to glycerolipids (green) and amino acids (blue). The peak at *m/z* 326.3781 (didecyldimethyl ammonium), is a commercial surface disinfectant unintentionally introduced to the tissue samples somewhere in the sample handling process.

### 2.3. Pro-inflammatory polymer implants significantly increased lipid levels in tissue near the implant

It is apparent that tissue glycerol lipids are elevated for the pro-inflammatory M1-polymer implant (Figure 3A). It has been demonstrated that glycerol lipid production influences immune cell activity and enhances inflammation (32). To probe the distribution of glycerol lipids as a function of distance from the implant in the subcutaneous adipose tissue, mass spectra were acquired from three different areas about 50 µm apart (next to the implant, mid-point and next to the dermis) as shown in Figure 3B, with glycerolipid intensity presented for each implant in Figures 3C-E and in Table S1, Supporting Information. The M1-polymer implant was associated with a higher intensity of glycerolipid peaks at the implant surface compared to silicone and the M2-polymer coated catheters (Figure 3D), thus indicating that glycerolipids are increased in pro-inflammatory tissue microenvironments. It is known that lipid molecules have potent immunologic effects that can affect inflammation and fibrosis (33, 34). A study by Schreib et al., reported that lipids deposited by host cell on the surface of a biomaterial after implantation, and the varieties of lipids deposited influence how the immune system reacts to the biomaterial. They noted that fibrotic PDMS implants exhibited greater quantities of fatty acids. The anti-inflammatory implants demonstrated a significant enrichment of anti-fibrotic phospholipids (35).

**Figure 3.**
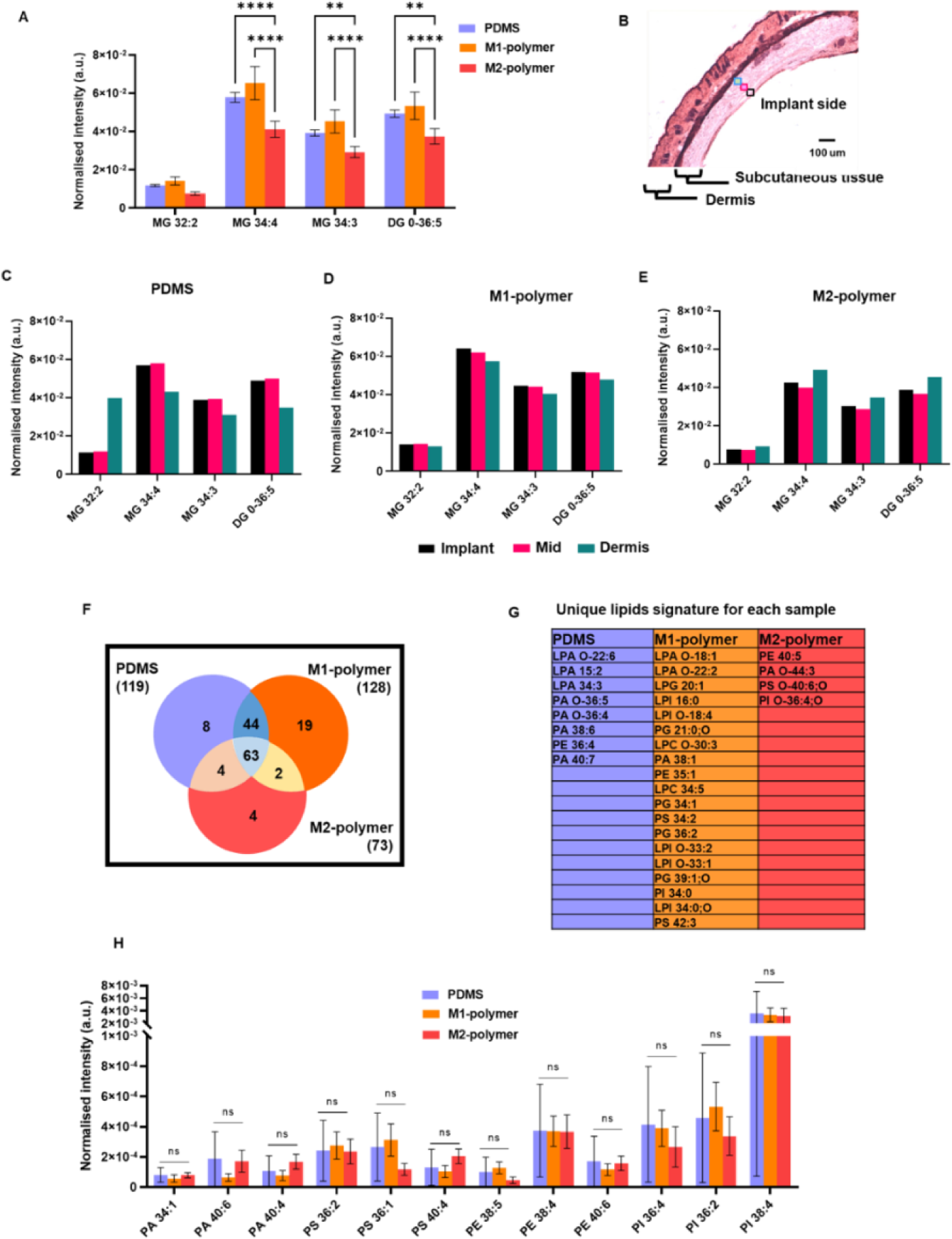
Relative quantification of characteristic lipid fragments detected by OrbiSIMS in positive and negative ion mode. (A) The normalized intensity of four different glycerolipid species in each tissue sample. (B) H&E stain image shows the three regions further away the implant was analyzed, implant (black), mid-point (pink) and dermis (sky blue). The normalized intensity of glycerolipid as a function of distance from the implant in each sample (C) PDMS, (D) M1-polymer and (E) M2-polymer. (F) Venn diagram comparison of the number of phospholipids detected in 3 different tissue samples using 3D OrbiSIMS and unique lipid signature for each sample. (G) The list of unique lipid signatures in each sample. (H) Normalized intensity of phospholipids in three separate samples.

Lipids are essential structural components of cell membranes, influencing membrane fluidity, ion exchange, and apoptotic signals and also major energy storage component (36). (37–40)A glycerolipid molecule combines glycerol and at least one fatty acid. Glycerolipids are the main long-term energy-storing molecules in mammalian cells and second messenger signaling lipid (41). Thus, much of the stored fat in the adipose tissues of animals consists primarily of glycerolipids (42). Monoacylglycerols (MG), diacylglycerols (DG) and triacylglycerols (TG) are types of glycerolipids (43). MG consists of a glycerol molecule connected to a single fatty acid. The two components are connected through an ester bond. DG is a glycerolipid composed of a glycerol molecule and two fatty acid chains linked by ester bonds. It has been demonstrated that glycerolipid production influence immune cell activity and enhances inflammation (44, 45). For example, DG functions as a second messenger that modulates the activation of protein kinase C (PKC), an enzyme that contributes in T-cell activation and proliferation, hence preserving the integrity of the cell membrane (46, 47). The accumulation of DG can lead to a state of lipotoxicity, which causes cell dysfunction and apoptosis, and can also induce diabetes and cancer (48). This work demonstrates that our tissue metabolite profiling with 3D OrbiSIMS successfully detected differential glycerolipid levels in tissue.

Phospholipid compounds were detected at lower ion intensities than the glycerol lipids, but those common and unique to the three tissues are compared in Figure 3F. A higher number of phospholipid compounds were detected from the tissue next to the M1 polymer than the uncoated silicone and the anti-inflammatory polymer. A total of 128 lipid peaks were putatively identified in tissue near the pro-inflammatory polymer, of which 63 lipid ions were common to the uncoated silicone and anti-inflammatory polymers. In addition, there were 4 unique lipid compounds in tissue near the anti-inflammatory polymer identified using the LIPID MAPS database (Table S2, Supporting Information). The list of unique phospholipid signatures in each sample are shown in Figure 3G. Almost all phospholipid species, phosphatidylserine (PS), phosphatidylethanolamine (PE) and phosphoinositides (PI) in tissue that saw M1 polymers were of a higher intensity than the M2 polymers (Figure 3H and Table S3, Supporting Information). However, a low signal from the PA species was also observed in the tissue sample from the pro-inflammatory polymer, and no statistically significant differences were observed between the implants. Recent research has focused on the various biological impacts of phosphatidic acid (PA) produced by activated macrophages and numerous other cells (38, 39). Particularly intriguing is the fact that PA functions as an intermediary messenger for several selective pro-inflammatory targets. PA has been reported to protect LPS-induced septic mice by pharmacologic inhibition (40).

The confirmation of the identity of some example putative lipid assignments was achieved using tandem mass spectrometry (MS/MS) in the Orbitrap when analyzing the tissue section sample. We performed MS/MS on tissue samples, confirming the negative ions assigned to the PA and PI as shown in the resulting spectra (Figure S4A-D, Supporting Information). Moreover, MS/MS confirmed the identity of the fragments from key mass ions in the tissue sample providing structural information on the key lipid species including fatty acid moiety, and lipid class.

### 2.4. Amino acid ion intensities are higher for M2 polymer implants

The *loadings* shown in Figure 2B indicated that the intensities of amino acid ions were strongly associated with the M2 polymer implant as quantified in Figure 4A, this is consistent with the collagenous layer preferentially around the M2-polymer implant (Figure S1C, Supporting Information). A total of 52 amino acid fragments were assigned from the tissue sample (Table S4, Supporting Information). Those differentiating between the samples are plotted in Figure 4A, with the M2 polymer implant exhibiting the highest intensities. Evaluating the ion intensity versus distance from the implant in Figure 4B-D, it is apparent that the greatest variance is seen in the tissue exposed to the M1 polymer implant, where the highest amino acid ion intensities are seen at the subcutaneous adipose tissue abutting the interface with the implant or at the dermis interface.

**Figure 4.**
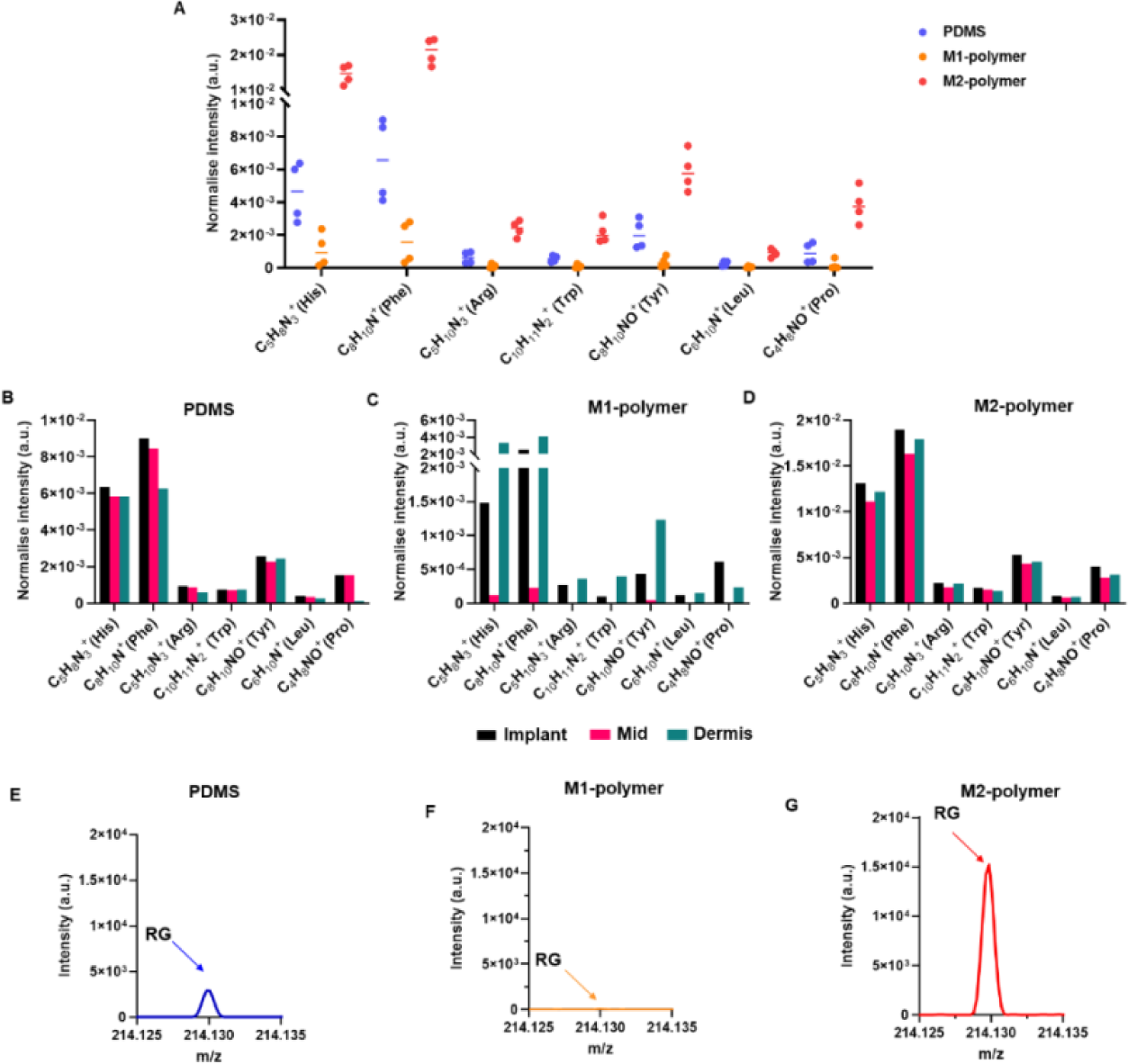
Characteristic amino acid fragments were detected in tissue section in positive ion mode. (A) The normalized intensity of amino acid fragments in each sample, PDMS (blue), M1-polymer (orange) and M2-polymer (red). The normalized intensity of amino acids with further away the implant in each sample (B) PDMS, C) M1-polymer and D) M2-polymer, implant (black), mid-point (pink) and dermis (sky blue). E-G) The spectrum of amino acids with RG sequences from each tissue sample.

The amino acid intensities are, more similar with distance for the uncoated silicone and M2 polymer implants (Figure 4C and D). The characteristic amino acid ions observed are consistent with fragmentation from proteins. Kotowska et al., (28) gathered lysozyme fragments in a spectrum from a protein monolayer sample, with the Arginine-Glycine (RG) amino acid pairs of lysozyme detected at *m/z* 214.1295 [C_8_H_16_N_5_O_2_]^+^. This protein fragment was also seen from the tissue samples and at similar relative intensities between samples, suggesting that these mono amino acid signals are fragments from proteins, and are not from free amino acids (Figure 4E-G and Table S5, Supporting Information).

### 2.5. Comparison of tissue and single cell metabolites

A range of non-lipid metabolites were detected in single cell macrophage expressed *in vitro* preferentially by M1, M2 or M0 cells that have been identified using the Human Metabolome Database and 3D OrbiSIMS (30). Pyridine (C_5_H_6_N^+^) and pyrimidine (C_4_H_5_N_2_^+^) had comparatively high ion intensities after M2 polarisation. This is consistent with our findings that pyridine moieties are intense in tissue adjacent to M2-polymer while being much lower in intensity for tissue adjacent to M1-polymer implants (Figure 5A and B). The single cell *in vitro* intensity versus the *in vivo* tissue intensity of pyridine and pyrimidine were compared and were shown to be highly correlative (simple linear regression curves: R^2^ = 0.91 for pyridine; R^2^ = 0.82 for pyrimidine), indicating that *in vitro* cell studies can semi-quantitatively predict *in vivo* pyridine and pyrimidine intensities.

**Figure 5.**
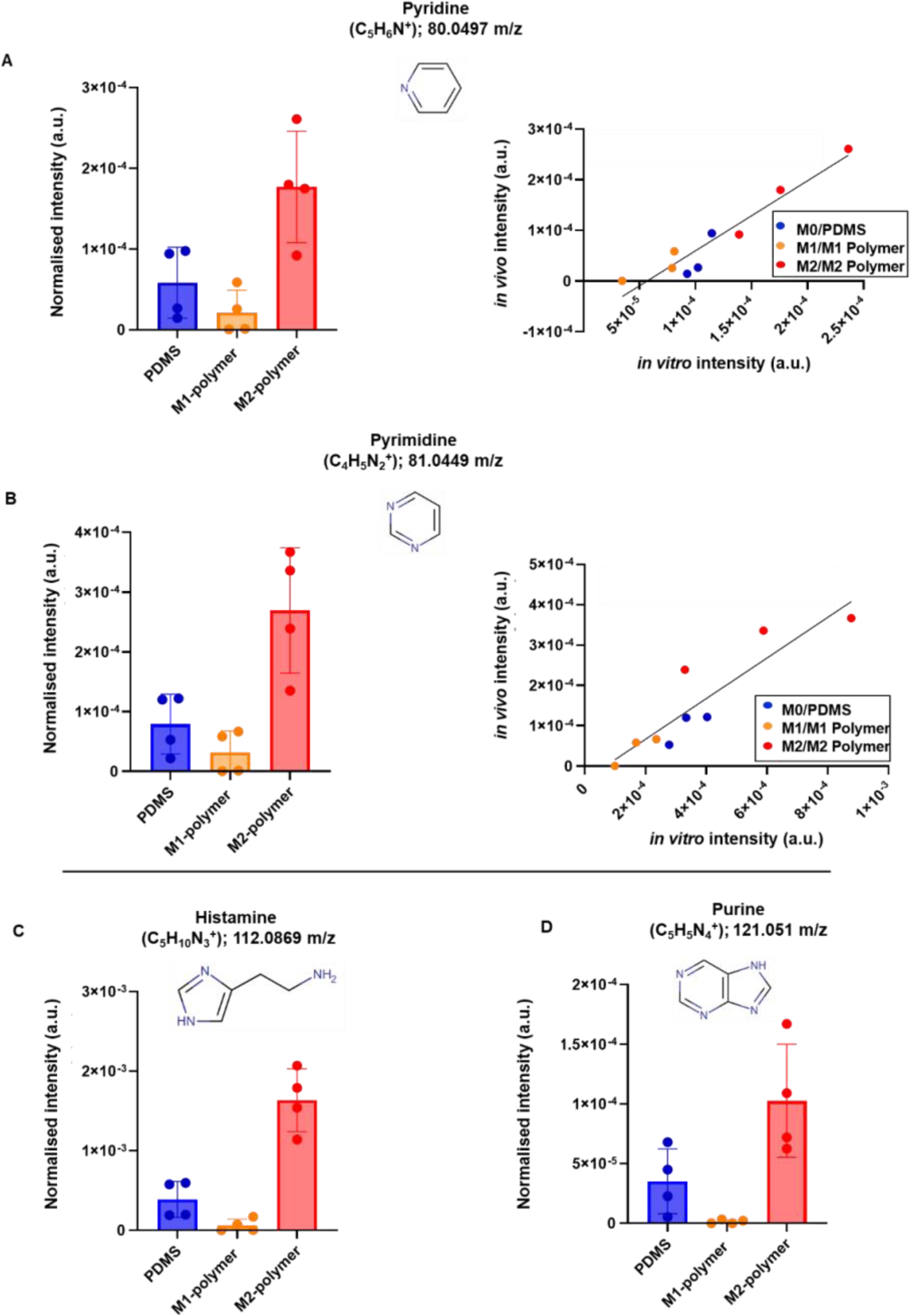
(A) Metabolites detected from *in vivo* tissue significantly affected by M1 and M2-polymers. (B) Comparing the compounds pyridine and pyrimidine from *in vivo* tissue samples and single cell macrophages analysis showing simple linear relationships for pyridine and for pyrimidine (52). (C) Histamine from *in vivo* and (D) Purine from *in vivo* are found predominantly in tissue adjacent to M2-polymer or PDMS.

### 2.6. Other small molecules found to be modulated by the implant coating

In tissue, histamine (C_5_H_10_N ^+^) and purine (C_5_H_5_N ^+^) were detected strongly from the M2 polymer tissue and not at all from M1 polymer (Figure 5C and D, Table S6, Supporting Information). Both compounds are connected to anti-inflammatory cellular reactions. Histamine can promote wound healing in skin lesions, inhibit tumour growth, and modulate inflammation in models of colitis and experimental autoimmune encephalomyelitis (EAE) (49). Purine, a common substrate in living organisms, is essential for cellular proliferation and a key regulator of the immune system. Multiple enzymes carefully regulate the purine de novo and salvage pathways, and malfunction in these enzymes results in excessive cell proliferation and immunological imbalance, which leads to tumour growth (50). Furthermore, purine has antioxidant and anti-inflammatory properties, as well as a role in cell energy homeostasis (51). These correlations between single cell analysis and tissue stimulated by implanted polymers gives support to the use of molecular characterisation to link *in vitro* and *in vivo* performance and its role in probing underpinning molecular mechanistic understanding.

### 2.7. Imaging of metabolites in the tissue abutting the implants

Using the 3D OrbiSIMS in imaging mode, the distribution of metabolites of interest from the above spectral analyses were produced as shown in Figure 6A-B from a 450 × 450 µm area of the subcutaneous adipose tissue interface adjacent to the implant. It is interesting that the M2 metabolites were located in bands near the interface between tissue and implant. As expected from the spectral analysis (Figure 5), they were seen more strongly in the M2-polymer image for pyridine, pyrimidine and histamine. The glycerides appear intense and uniformly distributed. (Figure 7A and B).

**Figure 6.**
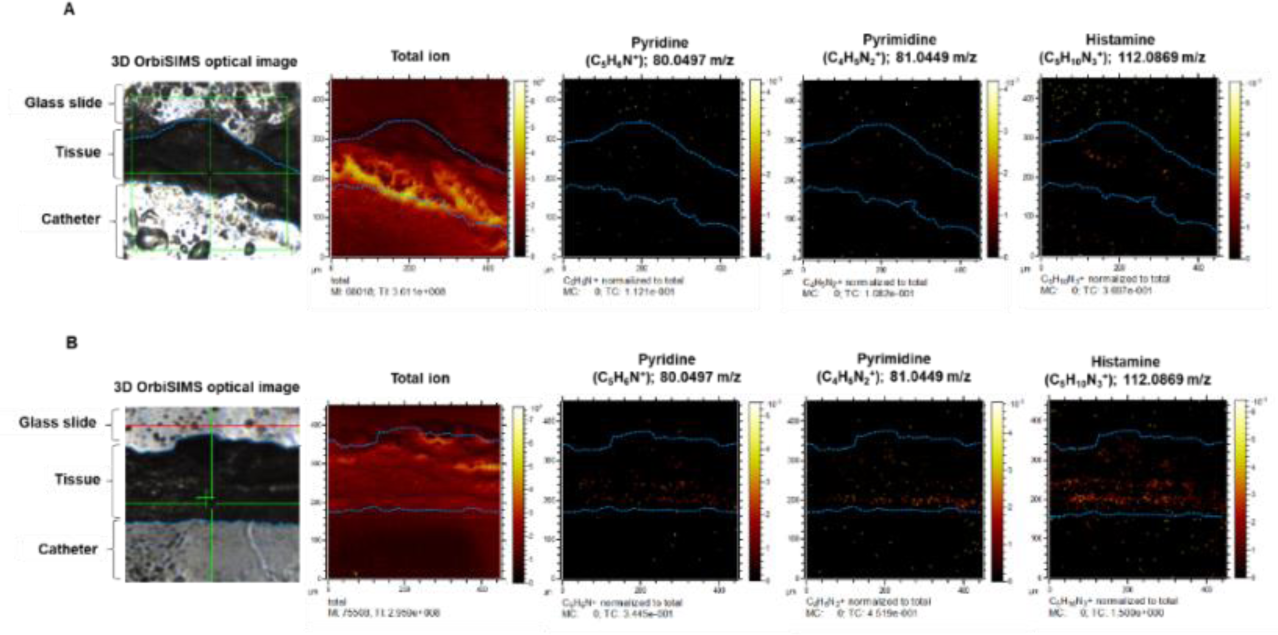
Chemical imaging of tissue sample. (A and B) Ion image of M1-plymer and M2-polymer were acquired (area of 450 µm × 450 µm), including pyridine, pyrimidine and histamine metabolite which are divided by total intensity.

**Figure 7.**
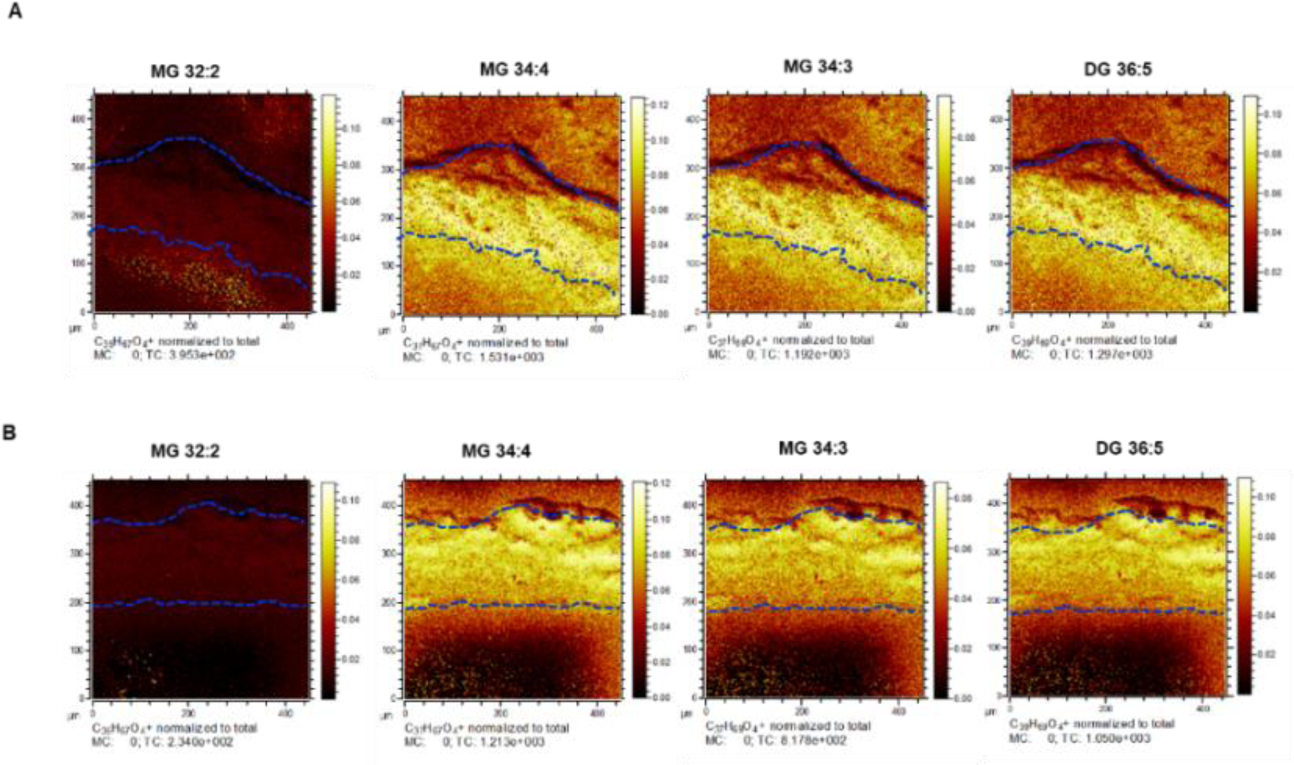
Chemical imaging of tissue sample. (A and B) Ion image of the sum of glycerolipids in M1-polymer and M2-polymer were acquired (area of 450 µm × 450 µm), including MG and DG which are divided by total intensity. The blue lines denoted the upper and lower boundaries of the of the tissue, with the catheter at the bottom of the image.

Representative molecular ions of the phospholipid classes PA, PS, PE and PI shown in Figure S3, Supporting Information. The PI ions were the most intense compared to other lipid species and the distribution of each ion seemed similar. In Figure S4, Supporting Information, we mapped the PI 38:4, [C_47_H_82_O_13_P]^-^ and [C_6_H_10_PO_8_]^-^ ion corresponding to the PI head group, and two fatty acids are represented by FA 18:0 [C_18_H_35_O_2_]^-^ peak at m/z 283.2642 and FA 20:0 [C_20_H_31_O_2_]^-^ peak at m/z 303.2327. We also mapped the nuclear marker and overlayed with the PI 34:4 marker as shown in Figure S3, Supporting Information. The ion contribution of nuclear marker [HP_2_O_6_]^-^ at m/z 158.9056 is mapped in blue, and PI 34:4 at m/z 885.5500 is mapped in red. The nuclear marker intensity distribution is more closely correlated to the phospholipids that the glycerides, suggesting that a proportion of these lower intensity former species are generated from cells.

## 3. Conclusion

We report a new label-free direct analysis strategy able to provide molecular insight into the host-implant interface using 3D OrbiSIMS. This study provides a detailed molecular characterization of tissue sections, allowing information on the distribution of metabolites, lipids and protein derived amino acids. Silicone catheter section coated with different immune-instructive polymers were used as example medical devices in a rodent model of FBR. Novel M1- and M2 inducing polymer coated implants produced distinct tissue metabolite profiles revealed by 3D OrbiSIMS. Interestingly, glycerolipids seem to be more abundant on pro-inflammatory surfaces. For some metabolites, the ion intensities were found to correlates with single cell analysis of polarised macrophages, highlighting the power of this approach in elucidating cell responses in complex biological context. Better understanding of immunometabolism has started to provide new insights into many pathologies as well as immune system’s role in maintaining tissue homeostasis. Combining powerful metabolic imaging techniques and biomaterials design could be transformative in enabling the design of novel bio-instructive materials that present positive interactions with the immune system to induce a pro-healing macrophage response following implantation.

## 4. Experimental Section

### Implant sample preparation

Clinical-grade silicone rubber catheters with a 13 mm external diameter were cut to a length of 5 mm (cylinder shape). To secure in an automated dip coating rig, microlance needles were inserted into the catheter wall and clamped. The catheters were prepared by dipping them into Nusil MED1-161 silicone primer, which is made up of tetrapropylsilicate and tetra (2-methoxyethoxy) in 50% v/v acetone and withdrawing rate of 1 mm/min for 30 seconds. They were then dried at room temperature for 2 min. MED1-161 coated catheters were dip-coated into a copolymer solution of each of the polymers (5 % w/v) in dichloromethane with a dipping and withdrawing rate of 1 mm/min for 30 seconds twice. Copolymers were synthesized via a thermal free radical polymerization method, purified by precipitation into an excess of methanol and then were characterized by NMR and GPC. The polymers used had previously been identified to polarize macrophages *in vitro* and modulate the FBR *in vivo*: Poly(cyclohexyl methacrylate-co-dimethylamino-ethyl methacrylate), pCHMA-DMAEMA (referred to as M1 polymer) which induce pro-inflammatory macrophage or poly(cyclohexyl methacrylate-co-isodecyl methacrylate), pCHMA-iDMA polymer (referred to as M2 polymer) which induce anti-inflammatory macrophage phenotype (12). Coated catheters were dried overnight at room temperature then dried in a vacuum at 50 °C (<0.3 mbar) for 7 days to remove solvent. The chemical structure of the monomers and copolymers pCHMA-co-DMAEMA and pCHMA-co-iDMA are presented in Table S7, Supporting Information.

### In vivo models

*In vivo* studies were approved by the University of Nottingham Animal Welfare and Ethical Review Board and carried out in accordance with Home Office authorization under project license number PP5768261. Female BALB/c mice, 19-22 g were used in these studies and were housed in individually ventilated cages (IVCs) under a 12 h light cycle, with access to food and water ad libitum. The weight and clinical condition of the mice were monitored daily. UV light was used for 20 minutes to sterilize the catheter segments prior to implantation. All segments were implanted subcutaneously into mice for 28 days, 3 mice/each polymer. At the end of the animal studies, on day 28, mice are humanely sacrificed by CO_2_ euthanasia. The polymer identity was blinded to the researchers until the end of the data quantification.

### Histological Analysis

The catheter segments and surrounding skin after 28 days of implantation were cut to approximately 5.5 cm × 5.5 cm and were embedded in optimal cutting temperature compound (OCT). Follow embedding, the tissue was placed in a cryostat chamber at -20°C and sliced into 15 µm thick sections (CM1850, Leica microsystems). The FBR to the polymer coatings was assessed by staining with haematoxylin and eosin (H&E) in Table S8, Supporting Information and Masson’s trichrome (MTC) using optimised protocols contained in Tables S9 and Table S10, Supporting Information respectively. Following H&E staining, images were recorded using an Axioplan microscope (Zeiss) with a 40X objective to count the number of macrophages and neutrophils (field of view 100 um x 100 um) N=2 and n=3. With MTC staining, each sample was captured at 10X magnification, and the thickness of the collagen measured by taking four measurements from the top to the bottom of the distinct layer: one at the top, one down and the other two further away cross of the images of each sample, N=3 and n=1.

### Macrophage phenotype analysis

The method used was taken from Rostam *et al*. (12). Briefly, tissue sections were stained to identify the macrophage phenotype at the catheter-tissue interface. This was carried out using the pro-inflammatory marker inducible nitric oxide synthase (iNOS) and the anti-inflammatory marker arginase-1 (Arg-1). The processing of the macrophage phenotype is shown in Table S4, Supporting Information (12). The stained cells were imaged with a Zeiss LSM880C confocal microscope, and any background fluorescence was subtracted using ImageJ. The mean raw fluorescence intensity density of the region of interest around the foreign body site was used to measure the sum of all pixels in the given area and at least five different fields of view were randomly examined in each tissue section. We used a signal-to-noise ratio (SNR) of 2 as the detection threshold for fluorescence intensity measurements by ImageJ software. The fluorescence intensity ratio of M2 macrophages to M1 macrophages was calculated for each region.

### OrbiSIMS analysis

After tissue sectioning, the slices of tissue supported on slides were gently washed 3 times with cold distilled water for 30 s, 1 time with cold 70% ethanol for 30 s and then 3 times with 1 mL of 150 mM ammonium formate solution for 30 s to remove salts which can decrease the sensitivity of molecules in SIMS by signal suppression (53). Tissue slides were plunged frozen in liquid nitrogen and then freeze dried for 12 hours to remove water whilst retaining some degree of 3D structure. The samples were subsequently stored in a microscope slide box container at -80 °C until analysis. Prior to OrbiSIMS analysis, the sample was warmed to room temperature without opening and then mounted onto the instrument sample holder and loaded into the 3D OrbiSIMS for analysis. 3D OrbiSIMS analysis was conducted with a Hybrid SIMS instrument (IONTOF, Germany) using Mode 4 which comprised single Ar ^+^ primary ion beam of energy of 20 keV a duty cycle of 4.4% and continuous GCIB current of 2.3 A, over an area of 100 × 100 µm with crater-size 180.0 × 180.0 µm collecting data using the OrbiTrap analyser in the mass range of *m/z* 75-1125. The electron flood gun was operated with an energy of 21 eV and an extraction bias of 20 V. for charge compensation. The pressure in the main chamber was maintained at 8.9 × 10^-7^ mbar using argon gas flooding. The OrbitrapTM cycle time was set to 200 µs. The Orbitrap analyzer was operated in positive and negative ion mode at the mass resolving power setting of 240,000 (at *m/z* 200). The secondary ion injection time was 500 ms, the total ion dose per measurement was 5.21 × 10^10^ ions/cm^2^. Adjacent areas on the tissue samples were analyzed, 4 regions surrounding the implant region (catheter-tissue interface) per one tissue section slide and 3 regions further away the implant (next to implant, mid-point and next to the dermis) were consumed with both positive and negative polarity.

One 3D OrbiSIMS image using the 20 keV Ar ^+^ analysis beam with a 2 µm diameter probe was acquired. The 20 µm analysis beam was configured as described in the spectra acquisition section. The pixel size 3 µm imaging beam duty cycle set to 37.7% and GCIB current was 2.3A. The images were run on the area of 450 × 450 µm using random raster mode. The cycle time was set to 400 μs. Argon gas flooding was in operation; to aid charge compensation, pressure in the main chamber was maintained at 9.0 × 10^−7^ mbar using argon gas flooding. The images were collected in negative polarity, in mass range 75–1125 *m/z*. The injection time was set to 500 ms, the total ion dose per measurement was 1.61 × 10^13^. Mass-resolving power was set to 240,000 at 200 *m/z*. All data analysis was carried out using Surface Lab 7.1 software (IONTOF GmbH).

### Principal component analysis (PCA)

The 3D OrbiSIMS spectra contained many secondary ions. Principal component analysis was applied to the data set to provide unbiased identification of the differences between each tissue sample. Spectra of all tissue samples were acquired by accumulating data from a single area, 4 areas of each sample were acquired, with each normalized to their respective total ion count in SurfaceLab 7 software. A peak list was constructed, containing the ions above the minimum ion count intensity, which was determined in each case as being greater than assigned noise signals (1428 peaks in the positive polarity spectra). Multivariate analysis of 3D OrbiSIMS results was done in simsMVA (https://mvatools.com/), using Matlab(54). The peak list was normalized to total ion count and applied to 4 regions of interest on all samples. The data was pre-processed by Poisson scaling and mean centring. PCA was run in algorithm mode, retaining all principal components.

### Data Processing and Metabolites Identification

Peak assignments for each sample were created by IonTOF SurfaceLab 7. Peak lists of secondary mass ions and secondary intensity from the software were exported and then imported into the LIPIDMAPS database to identify the lipid species. For metabolite results, 3D OrbiSIMS spectra were exported as .txt files then searched against the Human Metabolome Database with a 5 ppm mass tolerance for putative annotation.

## Supporting information

SuppInformation

## Supporting Information

Supporting Information is available from the Wiley Online Library or from the author.

## Acknowledgments

We acknowledge the financial support for this work from The Royal Thai Government Scholarship provided by the National Metal and Materials Technology Centre (MTEC), the National Science and Technology Development Agency (NSTDA), Thailand. This work was also supported by the Engineering and Physical Sciences Research Council (EPSRC) [grant number: EP/P029868/1] with a Strategic Equipment grant.

