## Supplementary material for "Spatially resolved molecular analysis of host response to medical device implantation using the 3D OrbiSIMS highlights a critical role for lipids": SuppInformation

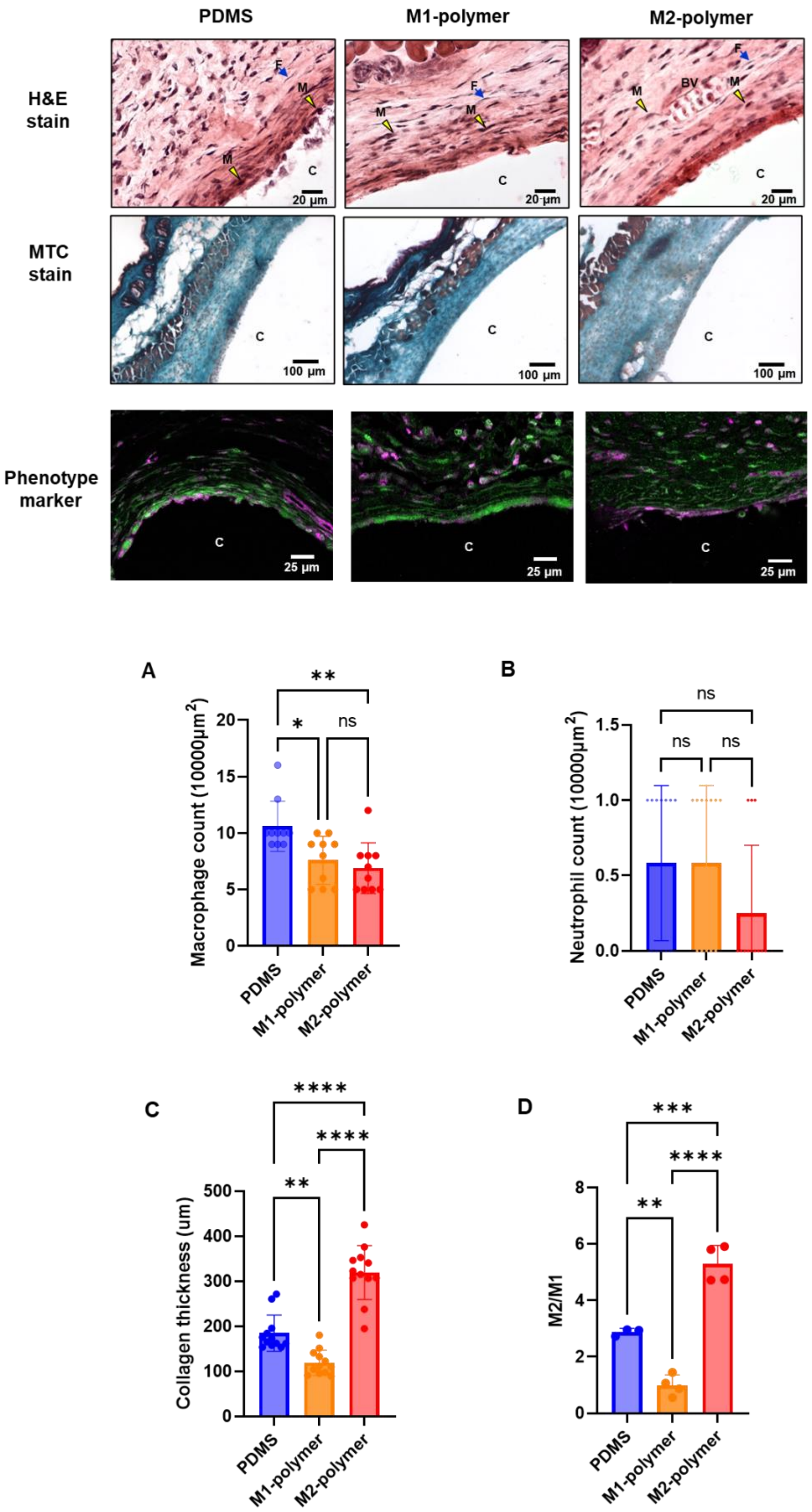

PDMS
M1-polymer
M2-polymer
H&E stain
MTC stain
Phenotype marker
F
M
BV
C
20 µm
100 µm
25 µm
A
Macrophage count (10000µm$^2$)
**
*
ns
B
Neutrophil count (10000µm$^2$)
ns
C
Collagen thickness (um)
****
**
D
M2/M1
***

**Fig S1.** Histological analysis of tissue sections following 28 days implantation of polymer coated catheter sections in a murine model of foreign body response. Representative H&E staining images, showing a well-defined inflammatory reaction fibroblast (F), blood vessels (BV), macrophage (M) and catheter (C) captured at × 40 magnification the scale bar = 20 µm. MTC staining image of each tissue section slide for identifying collagen thickness, captured at × 10 magnification. Scale bar = 100 µm. (A-C) show the infiltration count of macrophages, neutrophils and collagen thickness from the site surrounding the foreign body. All data are presented as the mean with ±s.d (N=2, n=3). Significance was calculated by one-way ANOVA with Tukey's post-hoc analysis: *$p<0.05$, ** $p<0.01$, *** $p<0.001$. (D) Representative images show tissue section stained for the M1 marker iNOS in green and M2 marker arginase1 in magenta and C represents the catheter site. (Images were acquired on confocal). Scale bar = 25 µm. (D) The ratio of M2-like macrophages to M1-like macrophages for each polymer. All data are presented as the mean with ±s.d (N = 2 and n = 5). Significance was calculated by one-way ANOVA with Tukey's post-hoc analysis: *** <0.0001

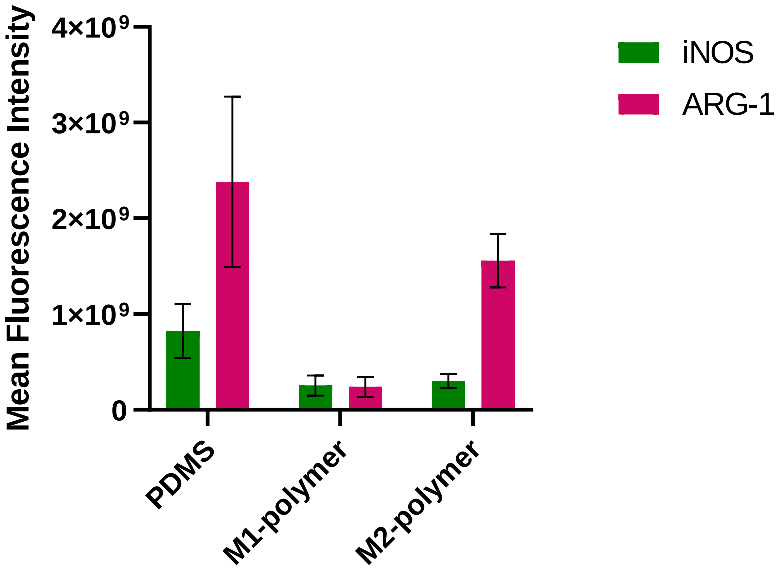


**Fig S2.** The mean fluorescescence intensity of iNOS and ARG-1 expression in tissue images. M2-polymer shown high level of the Arg-1 expression.

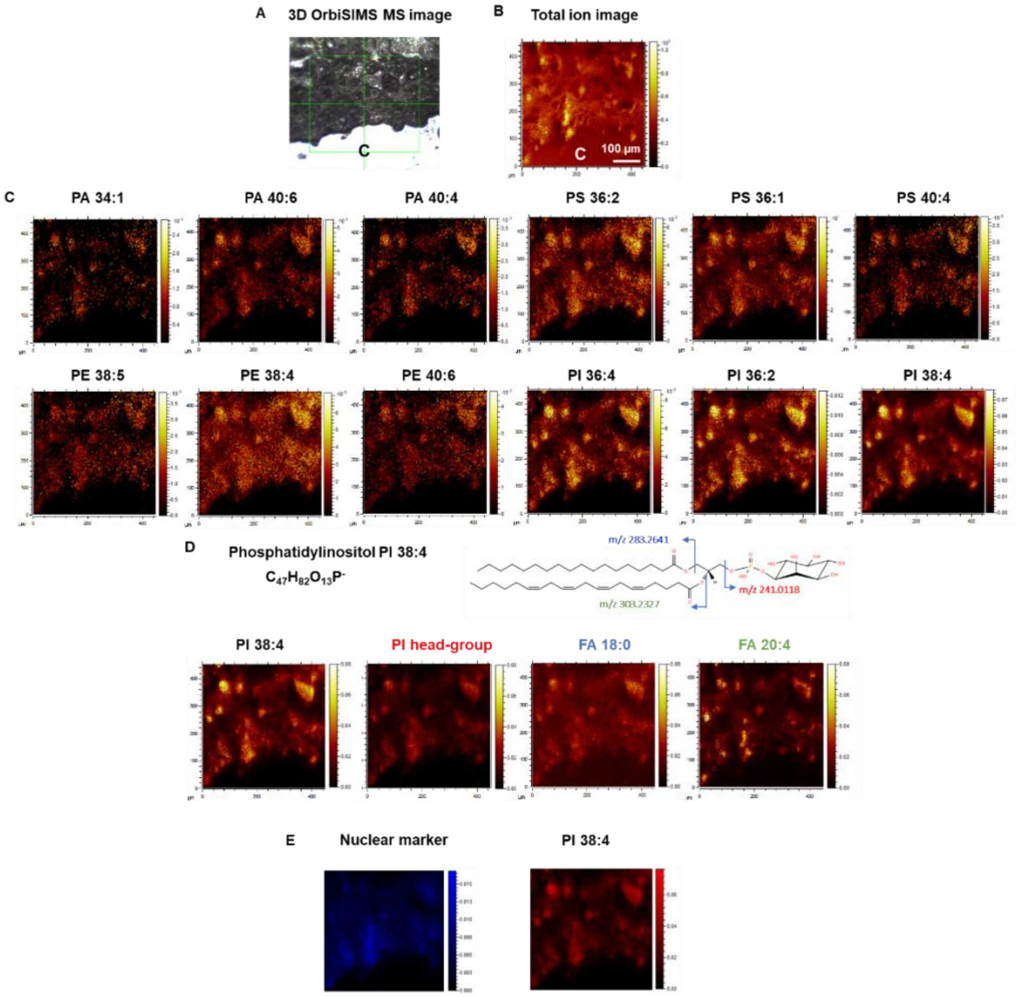


**Fig S3.** Chemical imaging of tissue sample. (A) View of the area where OrbiSIMS optical ion images were acquired (area of 450 μm × 450 μm). (B–D) 3D OrbiSIMS ion images were recorded in the negative ion mode, (B) total ion image, (C) ion image of the sum of phospholipid specie ions including PA, PS, PE and PI which are divided by total intensity. (D) The main ion of PI (38:4), showing the contribution of PI (38:4) ion are the signature fragments of PI head group, two fatty acids fragments (FA 18:0 and FA 20:4) and (E) RGB ion images showing nuclear marker (blue) and PA 34:4 (red).

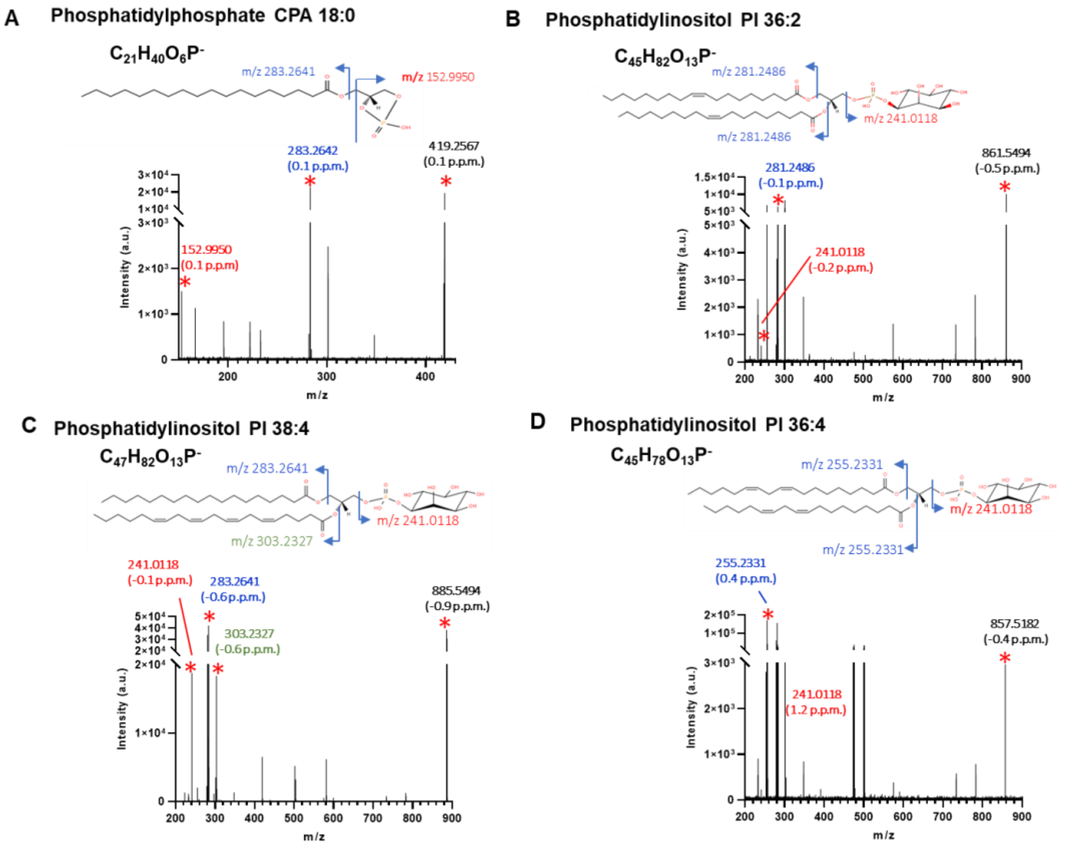


**Fig S4.** MS/MS product spectrum of lipids $[M\text{-}H]^-$. (A) The structure of CPA (18:0) at m/z 419.2517 was confirmed based on the detection of the PA head-group ions, $[C_6H_{10}PO_8]^-$ at m/z 241.0, and fatty acid moieties from these lipids are F18:0, $[C_{18}H_{35}O_2]^-$ at m/z 283.2642. (B) The precursor PI (36:2) ion is the signature fragments of PI headgroup, $[C_6H_{10}PO_8]^-$ and two FA18:1 fatty acid moieties are represented by $[C_{18}H_{33}O_2]^-$. and other MS/MS of PI (38:4) and (36:4) lipids are reported in Fig C and D)

**Table S1.** Glycerolipid fragments in 3D OrbiSIMS at surrounding implants in each sample PDMS, polymer M1-polymer and polymer M2-polymer. Monoglycerides (MG), diglycerides (DG)

| Mass m/z | Assignment | Glycerolipid | Average normalises Intensity (4 areas) | Average p.p.m (4 areas) |
|---|---|---|---|---|
| **PDMS** | | | | |
| 551.504 | $C_{35}H_{67}O_4^+$ | MG 32:2 | $1.18\times10^{-2}$ | 1 |
| 575.5038 | $C_{37}H_{67}O_4^+$ | MG 34:4 | $5.79\times10^{-2}$ | 0.6 |
| 577.5193 | $C_{37}H_{69}O_4^+$ | MG 34:3 | $3.92\times10^{-2}$ | 0.4 |
| 601.5195 | $C_{39}H_{69}O_4^+$ | DG 0-36:5 | $4.94\times10^{-2}$ | 0.6 |
| **M1-polymer** | | | | |
| 551.504 | $C_{35}H_{67}O_4^+$ | MG 32:2 | $1.42\times10^{-2}$ | 0.9 |
| 575.5038 | $C_{37}H_{67}O_4^+$ | MG 34:4 | $6.53\times10^{-2}$ | 0.8 |
| 577.5193 | $C_{37}H_{69}O_4^+$ | MG 34:3 | $4.53\times10^{-2}$ | 0.7 |
| 601.5195 | $C_{39}H_{69}O_4^+$ | DG 0-36:5 | $5.35\times10^{-2}$ | 0.8 |
| **M2-polymer** | | | | |
| 551.504 | $C_{35}H_{67}O_4^+$ | MG 32:2 | $7.50\times10^{-3}$ | 0.6 |
| 575.5038 | $C_{37}H_{67}O_4^+$ | MG 34:4 | $3.83\times10^{-2}$ | 0.3 |
| 577.5193 | $C_{37}H_{69}O_4^+$ | MG 34:3 | $2.93\times10^{-2}$ | 0.1 |
| 601.5195 | $C_{39}H_{69}O_4^+$ | DG 0-36:5 | $3.75\times10^{-2}$ | 0.3 |

**Table S2.** Show unique lipids signature for each phenotype; 8 unique for PDMS, 16 unique for M1-polymer and 4 unique for M2-polymer.

| Mass *m/z* | Formula $[M-H]^-$ | Name |
|---|---|---|
| **PDMS** | | |
| 467.2567 | $C_{25}H_{41}O_6P^-$ | LPA O-22:6 |
| 476.2784 | $C_{23}H_{44}NO_7P^-$ | LPA 15:2 |
| 655.4709 | $C_{37}H_{69}O_7P^-$ | LPA 34:3 |
| 679.4712 | $C_{39}H_{69}O_7P^-$ | PA O-36:5 |
| 681.4867 | $C_{39}H_{71}O_7P^-$ | PA O-36:4 |
| 719.4662 | $C_{41}H_{69}O_8P^-$ | PA 38:6 |
| 738.5083 | $C_{41}H_{74}NO_8P^-$ | PE 36:4 |
| 745.4819 | $C_{43}H_{71}O_8P^-$ | PA 40:7 |
| **M2-polymer** | | |
| 795.5743 | $C_{45}H_{85}NO_8P^-$ | LPI O-33:0 |
| 795.6271 | $C_{47}H_{89}O_7P^-$ | PA O-44:3 |
| 836.5446 | $C_{46}H_{80}NO_{10}P^-$ | PS O-40:6;O |
| 859.534 | $C_{45}H_{81}O_{13}P^-$ | PI O-36:4;O |
| **M1-polymer** | | |
| 421.2728 | $C_{21}H_{43}O_6P^-$ | LPA O-18:1 |
| 475.3195 | $C_{25}H_{49}O_6P^-$ | LPA O-22:2 |
| 537.32 | $C_{26}H_{51}O_9P^-$ | LPG 20:1 |
| 571.2891 | $C_{25}H_{49}O_{12}P^-$ | LPI 16:0 |
| 577.2787 | $C_{27}H_{47}O_{11}P^-$ | LPI O-18:4 |
| 585.3048 | $C_{26}H_{51}O_{12}P^-$ | PG 21:0;O |
| 670.5186 | $C_{38}H_{74}NO_6P^-$ | LPC O-30:3 |
| 729.5445 | $C_{41}H_{79}O_8P^-$ | PA 38:1 |
| 730.5395 | $C_{40}H_{78}NO_8P^-$ | PE 35:1 |
| 736.5294 | $C_{42}H_{76}NO_7P^-$ | LPC 34:5 |
| 747.5172 | $C_{40}H_{77}O_{10}P^-$ | PG 34:1 |
| 758.4981 | $C_{40}H_{74}NO_{10}P^-$ | PS 34:2 |
| 773.533 | $C_{42}H_{79}O_{10}P^-$ | PG 36:2 |
| 791.5433 | $C_{42}H_{81}O_{11}P^-$ | LPI O-33:2 |
| 793.559 | $C_{42}H_{83}O_{11}P^-$ | LPI O-33:1 |
| 833.5917 | $C_{45}H_{87}O_{11}P^-$ | PG 39:1;O |
| 837.5489 | $C_{43}H_{83}O_{13}P^-$ | PI 34:0 |
| 839.5643 | $C_{43}H_{85}O_{13}P^-$ | LPI 34:0;O |
| 868.608 | $C_{48}H_{88}NO_{10}P^-$ | PS 42:3 |

**Table S3**. Targeted phospholipid analysis in 3D OrbiSIMS spectra.

| Mass *m/z* | Assignment [M-H]$^-$ | Phospholipids | Area1 | | Area 2 | | Area 3 | | Area4 | |
|---|---|---|---|---|---|---|---|---|---|---|
| | | | Mass error p.p.m | Norm. intensity | Mass error p.p.m | Norm. intensity | Mass error p.p.m | Norm. intensity | Mass error p.p.m | Norm. intensity |
| **PDMS** | | | | | | | | | | |
| 673.4821 | $C_{37}H_{71}O_8P$ | PA 34:1 | 0.3 | $1.54\times10^{-4}$ | -0.1 | $5.52\times10^{-5}$ | 0.8 | $5.66\times10^{-05}$ | 1.0 | $5.79\times10^{-05}$ |
| 747.498 | $C_{43}H_{73}O_8P$ | PA 40:6 | 0.7 | $4.57\times10^{-4}$ | 0.3 | $7.92\times10^{-5}$ | 1.2 | $9.79\times10^{-05}$ | 0.8 | $1.10\times10^{-04}$ |
| 751.5291 | $C_{43}H_{77}O_8P$ | PA 40:4 | 0.3 | $2.57\times10^{-4}$ | 0 | $4.07\times10^{-5}$ | 0.9 | $5.80\times10^{-05}$ | 0.7 | $7.03\times10^{-05}$ |
| 786.5294 | $C_{42}H_{78}NO_{10}P$ | PS 36:2 | 0.4 | $5.42\times10^{-4}$ | -0.2 | $1.18\times10^{-4}$ | 0.9 | $1.40\times10^{-04}$ | 0.8 | $1.65\times10^{-04}$ |
| 788.5451 | $C_{42}H_{80}NO_{10}P$ | PS 36:1 | 0.5 | $6.03\times10^{-4}$ | -0.1 | $1.37\times10^{-4}$ | 1.3 | $1.55\times10^{-04}$ | 1.0 | $1.69\times10^{-04}$ |
| 838.5609 | $C_{46}H_{82}NO_{10}P$ | PS 40:4 | 0.6 | $3.10\times10^{-4}$ | 0.2 | $5.82\times10^{-5}$ | 1.2 | $7.27\times10^{-05}$ | 1.0 | $8.50\times10^{-05}$ |
| 764.5244 | $C_{43}H_{76}NO_8P$ | PE 38:5 | 0.4 | $2.47\times10^{-4}$ | 0 | $4.27\times10^{-5}$ | 1.1 | $5.37\times10^{-05}$ | 0.8 | $5.98\times10^{-05}$ |
| 766.5401 | $C_{43}H_{78}NO_8P$ | PE 38:4 | 0.4 | $8.32\times10^{-4}$ | 0.1 | $1.81\times10^{-4}$ | 1.1 | $2.31\times10^{-04}$ | 0.7 | $2.49\times10^{-04}$ |
| 790.5402 | $C_{45}H_{78}NO_8P$ | PE 40:6 | 0.4 | $4.20\times10^{-4}$ | 0 | $7.67\times10^{-5}$ | 0.9 | $8.64\times10^{-05}$ | 0.7 | $9.93\times10^{-05}$ |
| 857.5196 | $C_{45}H_{79}O_{13}P$ | PI 36:4 | 0.5 | $0.87\times10^{-4}$ | 0.2 | $1.79\times10^{-4}$ | 1.2 | $2.28\times10^{-04}$ | 1.0 | $2.66\times10^{-04}$ |
| 861.5511 | $C_{45}H_{83}O_{13}P$ | PI 36:2 | 0.5 | $1.10\times10^{-3}$ | 0.3 | $2.11\times10^{-4}$ | 1.3 | $2.37\times10^{-04}$ | 1.0 | $2.87\times10^{-04}$ |
| 885.5508 | $C_{47}H_{83}O_{13}P$ | PI 38:4 | 0.3 | $8.80\times10^{-3}$ | 0.2 | $1.54\times10^{-3}$ | 1.1 | $1.89\times10^{-03}$ | 0.9 | $2.04\times10^{-03}$ |
| **M1-polymer** | | | | | | | | | | |
| 673.4821 | $C_{37}H_{71}O_8P$ | PA 34:1 | 0.7 | $8.89\times10^{-05}$ | 0.3 | $2.82\times10^{-05}$ | 0.7 | $5.38\times10^{-05}$ | 0.6 | $5.68\times10^{-05}$ |
| 747.498 | $C_{43}H_{73}O_8P$ | PA 40:6 | 0.9 | $9.41\times10^{-05}$ | 0.9 | $3.33\times10^{-05}$ | 1.0 | $5.92\times10^{-05}$ | 1.1 | $6.99\times10^{-05}$ |
| 751.5291 | $C_{43}H_{77}O_8P$ | PA 40:4 | 0.7 | $1.23\times10^{-04}$ | 0.6 | $4.20\times10^{-05}$ | 0.6 | $6.79\times10^{-05}$ | 0.8 | $7.18\times10^{-05}$ |
| 786.5294 | $C_{42}H_{78}NO_{10}P$ | PS 36:2 | 0.7 | $3.88\times10^{-04}$ | 0.6 | $1.68\times10^{-04}$ | 0.9 | $2.59\times10^{-04}$ | 0.8 | $2.88\times10^{-04}$ |
| 788.5451 | $C_{42}H_{80}NO_{10}P$ | PS 36:1 | 0.7 | $4.45\times10^{-04}$ | 0.8 | $1.87\times10^{-04}$ | 1.0 | $2.91\times10^{-04}$ | 0.9 | $3.26\times10^{-04}$ |
| 838.5609 | $C_{46}H_{82}NO_{10}P$ | PS 40:4 | 1.0 | $1.55\times10^{-04}$ | 1.0 | $6.06\times10^{-05}$ | 1.1 | $9.17\times10^{-05}$ | 1.1 | $1.05\times10^{-04}$ |
| 764.5244 | $C_{43}H_{76}NO_8P$ | PE 38:5 | 0.6 | $1.64\times10^{-04}$ | 0.7 | $7.38\times10^{-05}$ | 0.9 | $1.26\times10^{-04}$ | 0.8 | $1.47\times10^{-04}$ |
| 766.5401 | $C_{43}H_{78}NO_8P$ | PE 38:4 | 0.6 | $4.80\times10^{-04}$ | 0.6 | $2.41\times10^{-04}$ | 0.9 | $3.54\times10^{-04}$ | 0.8 | $4.04\times10^{-04}$ |
| 790.5402 | $C_{45}H_{78}NO_8P$ | PE 40:6 | 0.5 | $1.64\times10^{-04}$ | 0.7 | $7.06\times10^{-05}$ | 0.7 | $1.07\times10^{-04}$ | 0.7 | $1.22\times10^{-04}$ |
| 857.5196 | $C_{45}H_{79}O_{13}P$ | PI 36:4 | 0.8 | $5.39\times10^{-04}$ | 0.7 | $2.49\times10^{-04}$ | 1.0 | $3.66\times10^{-04}$ | 1.1 | $4.02\times10^{-04}$ |
| 861.5511 | $C_{45}H_{83}O_{13}P$ | PI 36:2 | 0.8 | $7.32\times10^{-04}$ | 0.9 | $3.43\times10^{-04}$ | 1.0 | $5.06\times10^{-04}$ | 1.0 | $5.52\times10^{-04}$ |
| 885.5508 | $C_{47}H_{83}O_{13}P$ | PI 38:4 | 0.5 | $4.77\times10^{-03}$ | 0.6 | $2.11\times10^{-03}$ | 0.7 | $3.18\times10^{-03}$ | 0.7 | $3.42\times10^{-03}$ |
| **M2-polymer** | | | | | | | | | | |
| 673.4821 | $C_{37}H_{71}O_8P$ | PA 34:1 | 0.2 | $5.44\times10^{-05}$ | 0.2 | $8.07\times10^{-05}$ | 0 | $8.54\times10^{-05}$ | 0.3 | $9.54\times10^{-05}$ |
| 747.498 | $C_{43}H_{73}O_8P$ | PA 40:6 | 0.4 | $9.36\times10^{-05}$ | 0.6 | $1.54\times10^{-04}$ | 0.3 | $1.70\times10^{-04}$ | 0.6 | $2.70\times10^{-04}$ |
| 751.5291 | $C_{43}H_{77}O_8P$ | PA 40:4 | 0.1 | $1.04\times10^{-04}$ | 0.2 | $1.72\times10^{-04}$ | 0 | $1.77\times10^{-04}$ | 0.2 | $2.22\times10^{-04}$ |
| 786.5294 | $C_{42}H_{78}NO_{10}P$ | PS 36:2 | 0.1 | $1.36\times10^{-04}$ | 0.3 | $2.39\times10^{-04}$ | 0.2 | $2.32\times10^{-04}$ | 0.3 | $3.36\times10^{-04}$ |

| | | | | | | | | | | |
|---|---|---|---|---|---|---|---|---|---|---|
| 788.5451 | $C_{42}H_{80}NO_{10}P$ | PS 36:1 | 0.3 | $7.07\times10^{-05}$ | 0.5 | $1.26\times10^{-04}$ | 0.3 | $1.11\times10^{-04}$ | 0.4 | $1.65\times10^{-04}$ |
| 838.5609 | $C_{46}H_{82}NO_{10}P$ | PS 40:4 | 0.3 | $1.42\times10^{-04}$ | 0.4 | $2.05\times10^{-04}$ | 0.4 | $2.08\times10^{-04}$ | 0.5 | $2.61\times10^{-04}$ |
| 764.5244 | $C_{43}H_{76}NO_8P$ | PE 38:5 | 0.2 | $2.29\times10^{-05}$ | 0.1 | $4.06\times10^{-05}$ | 0 | $4.36\times10^{-05}$ | 0.2 | $7.48\times10^{-05}$ |
| 766.5401 | $C_{43}H_{78}NO_8P$ | PE 38:4 | 0.3 | $2.31\times10^{-04}$ | 0.3 | $3.72\times10^{-04}$ | 0.1 | $3.65\times10^{-04}$ | 0.3 | $5.03\times10^{-04}$ |
| 790.5402 | $C_{45}H_{78}NO_8P$ | PE 40:6 | 0.3 | $9.42\times10^{-05}$ | 0.3 | $1.62\times10^{-04}$ | 0.1 | $1.65\times10^{-04}$ | 0.3 | $2.10\times10^{-04}$ |
| 857.5196 | $C_{45}H_{79}O_{13}P$ | PI 36:4 | 0.3 | $1.33\times10^{-04}$ | 0.3 | $2.31\times10^{-04}$ | 0.1 | $2.52\times10^{-04}$ | 0.4 | $4.50\times10^{-04}$ |
| 861.5511 | $C_{45}H_{83}O_{13}P$ | PI 36:2 | 0.4 | $1.80\times10^{-04}$ | 0.4 | $3.33\times10^{-04}$ | 0.2 | $3.45\times10^{-04}$ | 0.5 | $4.93\times10^{-04}$ |
| 885.5508 | $C_{47}H_{83}O_{13}P$ | PI 38:4 | 0.4 | $1.76\times10^{-03}$ | 0.3 | $2.97\times10^{-03}$ | 0.1 | $3.04\times10^{-03}$ | 0.3 | $4.78\times10^{-03}$ |

**Table 4.** Characteristic molecular ion and fragments of amino acid in 3D OrbiSIMS spectra (positive polarity)

| Mass m/z | Assignment | Amino acids |
|---|---|---|
| 80.0498 | $C_5H_6N^+$ | Leucine |
| 86.0967 | $C_5H_{12}N^+$ | Isoleucine |
| 81.045 | $C_4H_5N_2^+$ | Histidine |
| 82.0528 | $C_4H_6N_2^+$ | Histidine |
| 93.0449 | $C_5H_5N_2^+$ | Histidine |
| 94.0527 | $C_5H_6N_2^+$ | Histidine |
| 95.0605 | $C_5H_7N_2^+$ | Histidine |
| 110.0713 | $C_5H_8N_3^+$ | Histidine |
| 156.0768 | $C_6H_{10}N_3O_2^+$ | Histidine |
| 100.087 | $C_4H_{10}N_3^+$ | Arginine |
| 112.0869 | $C_5H_{10}N_3^+$ | Arginine |
| 114.1026 | $C_5H_{12}N_3^+$ | Arginine |
| 120.0444 | $C_7H_6NO^+$ | Tryptophan |
| 130.0652 | $C_9H_8N^+$ | Tryptophan |
| 131.073 | $C_9H_9N^+$ | Tryptophan |
| 132.0808 | $C_9H_{10}N^+$ | Tryptophan |
| 143.073 | $C_{10}H_9N^+$ | Tryptophan |
| 157.0761 | $C_{10}H_9N_2^+$ | Tryptophan |
| 158.0839 | $C_{10}H_{10}N_2^+$ | Tryptophan |
| 159.0917 | $C_{10}H_{11}N_2^+$ | Tryptophan |
| 84.0447 | $C_4H_6NO^+$ | Glutamic acid |
| 84.0811 | $C_5H_{10}N^+$ | Lysine |
| 86.0603 | $C_4H_8NO^+$ | Hydroxyproline |
| 87.0555 | $C_3H_7N_2O^+$ | Asparagine |
| 120.0808 | $C_8H_{10}N^+$ | Phenylalanine |
| 166.0863 | $C_9H_{12}NO_2^+$ | Phenylalanine |
| 101.071 | $C_4H_9N_2O^+$ | Glutamine |
| 130.0499 | $C_5H_8NO_3^+$ | Glutamine |
| 136.0758 | $C_8H_{10}NO^+$ | Tyrosine |
| 104.053 | $C_4H_{10}NS^+$ | Methionine |
| 116.0706 | $C_5H_{10}NO_2^+$ | Proline |
| 82.0654 | $C_5H_8N^+$ | Multiple amino acids |
| 83.0607 | $C_4H_7N_2^+$ | Multiple amino acids |
| 88.0396 | $C_3H_6NO_2^+$ | Multiple amino acids |
| 96.0809 | $C_6H_{10}N^+$ | Multiple amino acids |
| 98.0966 | $C_6H_{12}N^+$ | Multiple amino acids |
| 100.0394 | $C_4H_6NO_2^+$ | Multiple amino acids |
| 102.055 | $C_4H_8NO_2^+$ | Multiple amino acids |
| 107.0492 | $C_7H_7O^+$ | Multiple amino acids |
| 114.055 | $C_5H_8NO_2^+$ | Multiple amino acids |
| 117.0573 | $C_8H_7N^+$ | Multiple amino acids |
| 118.0651 | $C_8H_8N^+$ | Multiple amino acids |
| 119.0492 | $C_8H_7O^+$ | Multiple amino acids |
| 128.0706 | $C_6H_{10}NO_2^+$ | Multiple amino acids |
| 121.0648 | $C_8H_9O^+$ | Multiple amino acids |
| 77.0389 | $C_6H_5^+$ | Generic fragment |
| 80.0624 | $C_6H_8^+$ | Generic fragment |
| 89.0388 | $C_7H_5^+$ | Generic fragment |
| 91.0545 | $C_7H_7^+$ | Generic fragment |
| 102.0465 | $C_8H_6^+$ | Generic fragment |
| 103.0543 | $C_8H_7^+$ | Generic fragment |
| 105.0699 | $C_8H_9^+$ | Generic fragment |

**Table 5.** Peak exported from SurfaceLab positive mode from each tissue sample, consisting of ions detected in the spectrum and assigned as RG sequences of lysozyme $[M-H]^+$ $C_8H_{16}N_5O_2^+$, *m/z* 124.1298

| **Sample** | **Area1** | | **Area 2** | | **Area 3** | | **Area4** | |
|---|---|---|---|---|---|---|---|---|
| | **Mass error p.p.m** | **Norm. intensity** | **Mass error p.p.m** | **Norm. intensity** | **Mass error p.p.m** | **Norm. intensity** | **Mass error p.p.m** | **Norm. intensity** |
| **PDMS** | 0.4 | $2.51\times10^{-05}$ | 0.6 | $4.09\times10^{-06}$ | -0.1 | $2.30\times10^{-06}$ | 0.2 | $2.67\times10^{-05}$ |
| **M1-polymer** | 0.2 | $1.42\times10^{-06}$ | 0.3 | $5.59\times10^{-05}$ | 8.7 | 0 | -6 | 0 |
| **M2-polymer** | 0.2 | $1.03\times10^{-04}$ | -0.3 | $8.89\times10^{-05}$ | -0.1 | $1.03\times10^{-04}$ | 0.2 | $1.43\times10^{-04}$ |

**Table 6.** Other small molecules in each sample and search by human metabolome data base.

| Mass *m/z* | Assignment $[M-H]^-$ | Metabolites | Area1 | | Area 2 | | Area 3 | | Area4 | |
|---|---|---|---|---|---|---|---|---|---|---|
| | | | Mass error p.p.m | Norm. intensity | Mass error p.p.m | Norm. intensity | Mass error p.p.m | Norm. intensity | Mass error p.p.m | Norm. intensity |
| **PDMS** | | | | | | | | | | |
| 80.0497 | $C_5H_6N^+$ | Pyridine | 4.1 | $9.77\times10^{-05}$ | 4.1 | $1.46\times10^{-05}$ | 3.9 | $2.67\times10^{-05}$ | 3.7 | $9.44\times10^{-05}$ |
| 81.0449 | $C_4H_5N_2^+$ | Pyrimidine | 3.9 | $1.20\times10^{-04}$ | 4.2 | $2.19\times10^{-05}$ | 3.9 | $5.30\times10^{-05}$ | 3.6 | $1.22\times10^{-05}$ |
| 112.0869 | $C_5H_{10}N_3^+$ | Histamine | -0.1 | $5.95\times10^{-04}$ | -0.2 | $1.90\times10^{-04}$ | -0.2 | $1.96\times10^{-04}$ | -0.4 | $5.76\times10^{-04}$ |
| 121.051 | $C_5H_5N_4^+$ | Purine | 0.2 | $4.49\times10^{-05}$ | 0.4 | $5.31\times10^{-06}$ | 0.0 | $2.26\times10^{-05}$ | -0.1 | $6.79\times10^{-05}$ |
| **M1-polymer** | | | | | | | | | | |
| 80.0497 | $C_5H_6N^+$ | Pyridine | 4.5 | $5.89\times10^{-05}$ | 4.5 | $2.57\times10^{-05}$ | 4.0 | $5.71\times10^{-05}$ | 3.5 | $1.08\times10^{-05}$ |
| 81.0449 | $C_4H_5N_2^+$ | Pyrimidine | 4.0 | $5.87\times10^{-05}$ | 3.7 | $6.67\times10^{-05}$ | 4.1 | $5.03\times10^{-05}$ | 3.9 | $1.27\times10^{-05}$ |
| 112.0869 | $C_5H_{10}N_3^+$ | Histamine | 0.0 | $1.70\times10^{-04}$ | -0.1 | $7.32\times10^{-05}$ | 4.2 | $6.78\times10^{-05}$ | 4.0 | $3.03\times10^{-05}$ |
| 121.051 | $C_5H_5N_4^+$ | Purine | -0.1 | $2.13\times10^{-06}$ | 0.2 | $3.34\times10^{-06}$ | - | 0 | - | 0 |
| **M2-polymer** | | | | | | | | | | |
| 80.0497 | $C_5H_6N^+$ | Pyridine | 3.4 | $1.75\times10^{-04}$ | 3.2 | $9.20\times10^{-05}$ | 0 | $1.80\times10^{-04}$ | 0.3 | $2.61\times10^{-04}$ |
| 81.0449 | $C_4H_5N_2^+$ | Pyrimidine | 3.1 | $2.39\times10^{-05}$ | 3.2 | $1.35\times10^{-04}$ | 3.3 | $3.36\times10^{-04}$ | 3.3 | $3.67\times10^{-04}$ |
| 112.0869 | $C_5H_{10}N_3^+$ | Histamine | -0.9 | $1.54\times10^{-03}$ | -0.9 | $1.14\times10^{-03}$ | -0.7 | $1.79\times10^{-03}$ | -0.8 | $2.07\times10^{-03}$ |
| 121.051 | $C_5H_5N_4^+$ | Purine | -0.5 | $6.25\times10^{-05}$ | -0.6 | $7.19\times10^{-05}$ | -0.4 | $1.67\times10^{-04}$ | -0.4 | $1.09\times10^{-04}$ |

**Table S7.** Chemical structure of the monomers and synthesis of copolymers, CHMA-co-DMAEMA and CHMA-co-iDMA.

| **Code** | **Monomer 1 name/structure (66%)** | **Monomer 2 name/structure (33%)** | **Copolymers** |
|---|---|---|---|
| M1-polymer | 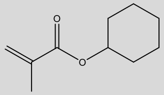 Cyclohexyl methacrylate (CHMA) | 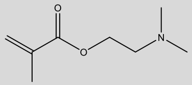 Dimethylamino-ethylmethacrylate (DMAEMA) | 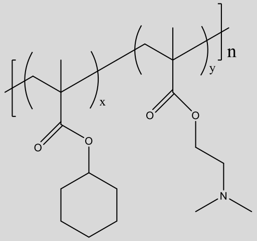 CHMA-co-DMAEMA |
| M2-polymer | 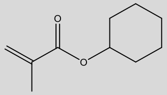 Cyclohexyl methacrylate (CHMA) | 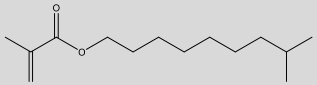 Isodecyl methacrylate (iDMA) | 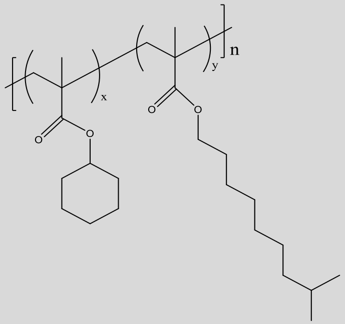 CHMA-co-iDMA |

**Table S2** Hematoxylin and Eosin (H&E) staining schedule.

| | | |
|---|---|---|
| 1. | Water rinse to remove OCT | 5 min |
| 2. | Haematoxylin solution | 5 min |
| 3. | Water rinse | 1 min |
| 4. | 1 % acetic acid in alcohol | 30 s |
| 5. | Water rinse | 1 min |
| 6. | Alkine Scott's | 1 min |
| 7. | Water rinse | 1 min |
| 8. | Eosin | 2 min |
| 9. | Water rinse | 1 min |
| 10. | 50 % alcohol | 30 s |
| 11. | 70 % alcohol | 30 s |
| 12. | 90 % alcohol | 30 s |
| 13. | 100 % alcohol × 2 | 30 s each |
| 14. | Xylene × 2 | 2 min each |
| 15. | Mounting media onto tissue slide and covered with a thin coverslip | |

**Table S3.** Masson Trichrome Stain Kit (Light Green) Masson 1929 schedule

| | | |
|---|---|---|
| 1. | Fixing, 4 %PFA | 1 h |
| 2. | Water rinse | 5 min |
| 3. | Haematoxylin, mixing equal volumes of Weigerts solution A & B (1:1) as required | 20 min |
| 4. | Water rinse | 1 min |
| 5. | 1 % acetic acid in alcohol | 30 s |
| 6. | Water rinse | 1 min |
| 7. | Ponceau fuchsin Masson solution for | 5 min |
| 8. | Rinse in distilled water | 2 min |
| 9. | The light green solution | 3 min |
| 10. | Water rinse | 30 s |
| 11. | 50 % alcohol | 30 s |
| 12. | 70 % alcohol | 30 s |
| 13. | 90 % alcohol | 30 s |
| 14. | 100 % alcohol × 2 | 30 s each |
| 15 | Xylene × 2 | 2 min each |
| 16. | Mounting media onto tissue slide and covered with a thin coverslip | |

**Table S4.** Sequential antibody staining for macrophage marker schedule

| | | |
|---|---|---|
| 1. | Washing in 0.2% Tween 20 in PBS × 3 | 5 min |
| 2. | 0.1% Triton X-100 in PBS | 10 min |
| 3. | Washing in 0.2% Tween 20 in PBS × 3 | 5 min |
| 4. | 5% BSA and plus 5% donkey serum in PBS | 1 h |
| 5. | 0.2% PBS-Tween 20 rinse × 3 | 5 min |
| 6. | Add diluted primary antibody with 1:50 of rabbit anti-mouse iNOS (Abcam) and 1:50 of goat anti-mouse Arg-1 (Thermo Fisher Scientific) in 5% goat serum at 4ºC | Overnight |
| 7. | Washing in 0.2% Tween 20 in PBS × 3 | 5 min |
| 8. | Add diluted secondary antibodies, donkey anti-goat IgG (H + L), and donkey anti-rabbit IgG (H + L) labelled with Alexa Fluor-594 and -488 (1:200; A11058 and A21206, Thermo Fisher Scientific), | 1 h |
| 9. | Washing in 0.2% Tween 20 in PBS × 3 | 5 min |
| 10. | 4',6 Diamidino-2-Phenylindole (DAPI, 20000 ng/ml) | 5 min |
| 11. | Washing in 0.2% Tween 20 in PBS × 2 | 5 min |
| 12. | Final, washing in distilled water | 5 min |
| 13. | Mounting media onto tissue slide and covered with a thin coverslip | |
